# Dynamic fingerprinting of the human functional connectome

**DOI:** 10.1101/2025.02.20.637919

**Authors:** Amin Ghaffari, Yufei Zhao, Xu Chen, Jason Langley, Xiaoping Hu

**Author notes:** Correspondence to: Xiaoping P. Hu, Ph.D., University of California, Riverside, Department of Bioengineering, Materials Science and Engineering 205.

## Abstract

Resting-state functional connectivity (FC) have distinct, personalized patterns that could serve as a unique fingerprint of each individual’s brain. While previous brain fingerprinting methods have used functional connectivity maps over a scanning session (static method), it has been shown that the brain is a dynamic system that switches between several metastable states, each of which having a different FC map. Taking the dynamic nature of brain connectivity into account will likely lead to more subject-specific information and better individual identification. In this paper, we derived the state-specific FCs using sliding window correlation and clustering and evaluated their performance in individual identification and cognitive score prediction. The resultant dynamic fingerprints outperformed the static fingerprints in identification accuracy. Furthermore, some of the brain states were more accurate in predicting cognitive scores, indicating that connectivity in some brain states is informative of cognition abilities, possibly useful as biomarkers for brain disorders.

**Impact Statement:** Our findings suggest that state-specific functional connectivity patterns of the brain are unique for each individual. These brain states predicted participants’ cognitive performance, suggesting that they have the potential to be used as biomarkers for cognitive function or neurological disorders. Integrating state-based functional connectivity into clinical frameworks could potentially enhance early diagnosis and patient stratification, and lead to targeted interventions for neuropsychiatric and neurodegenerative conditions.

## 1 Introduction

There is a uniqueness in the functional architecture of the brain that is specific to each individual (Airan et al., 2016, Gao et al., 2014, Mueller et al., 2013). This uniqueness is reflected in the functional connectivity map of the brain (i.e. connectome) derived from resting-state functional MRI (rs-fMRI), which encodes individual-specific information about cognition and behavior (Baldassarre et al., 2012, Cui and Gong, 2018, Jaywant et al., 2023, Shen et al., 2017), attention (Rosenberg et al., 2018), cognitive disorders (Panda et al., 2022, Sankar et al., 2023, Schiff, 2023, Tewarie et al., 2015, Wang et al., 2021b), and demographic factors (Panda et al., 2014, Smith et al., 2015). Given the diverse individual-specific information it provides, functional connectivity offers valuable potential for identifying markers of intelligence, behavior, and neurodegenerative disorders.

Earlier work examining connectome individuality used a stationary approach and found that functional connectome can serve as a fingerprint to distinguish different individuals (Finn et al., 2015). While considering the stationary functional connectivity maps as brain fingerprints is a well-established concept, researchers have also improved the performance of fingerprinting by applying advanced techniques on the stationary connectome such as group-wise decomposition (Amico and Goñi, 2018) and autoencoders (Cai et al., 2021).

The aforementioned static rs-fMRI studies assumed that the functional connectome is temporally invariant, but a great deal of studies revealed that the rs-fMRI functional connectome is organized into metastable states that change over time (de Vos et al., 2018, Greene et al., 2023, Liegeois et al., 2017). Static rs-fMRI analyses generate functional connectome by computing the correlation coefficient between different pairs of brain regions for the entire time series, and ignore the dynamic information (Allen et al., 2014, Calhoun and Adali, 2016), thereby leaving out any individual variability that exists in the dynamics of brain states (Liu et al., 2018). Incorporating temporal information of brain connectivity has improved identification accuracy of fingerprints as compared to the static method (Wang et al., 2019).

In this study, we used a sliding time-window-based approach along with clustering to extract state-specific functional connectomes (Chang et al., 2013, Handwerker et al., 2012) and developed a dynamic FC fingerprinting framework. With this approach, we could find each participant’s connectome in each metastable brain state, and extract individual-based information out of them. We hypothesized that each different brain state provides specific information contributing to individual identification, and aggregation of them is a more robust method for fingerprinting. Furthermore, we compared identification accuracy of our dynamic FC fingerprinting approach with the static fingerprinting approach and predicted the participants’ performance of cognitive tasks and found the key resting-state networks associated with these tasks. Our approach allows us to capture the dynamic nature of brain connectivity and leads to more accurate predictions of cognitive performance, potentially identifying better biomarkers for various neurological conditions.

## 2 Methods

### 2.1 Dataset and preprocessing

Resting-state data of 220 healthy participants (M_age_=29.2 yrs, age range=22-36; 125 females) were taken from the Q1-Q3 releases of the Human Connectome Project (HCP) S500 dataset (Van Essen et al., 2012) and used in our fingerprinting analysis. The resting-state scans for these participants were collected in separate sessions on two different days, with each session consisting of two 14.4-minute (1200-volume) acquisitions with opposite phase encoding directions. Full details about the scanning procedure and parameters can be found in related literature (Van Essen et al., 2013). Preprocessing of raw images was performed according to the HCP minimal preprocessing pipeline which includes removal of gradient distortion artifacts, correcting for motion and image distortion, registering to the Montreal Neurological Institute (MNI) space, and normalization (Glasser et al., 2013). The Institutional Review Board of Washington University in St. Louis approved the study, and all participants gave written consent before the first scan session. Resting-state data of five participants were dropped due to incomplete data (less acquired volumes), resulting in 215 participants for this analysis.

### 2.2 Parcellation and window-based connectomes

The Shen et al. 268-node parcellation (Figure 1.a) was used to define the regions of interest (ROIs) for the fingerprinting analysis in the MNI space (Shen et al., 2013). The regions in this parcellation are grouped into eight functional networks, which are the medial frontal network, the frontoparietal network, the default mode network, the subcortical-cerebellum network, the motor network, the visual I network, the visual II network, and the visual association network.

**Figure 1.**
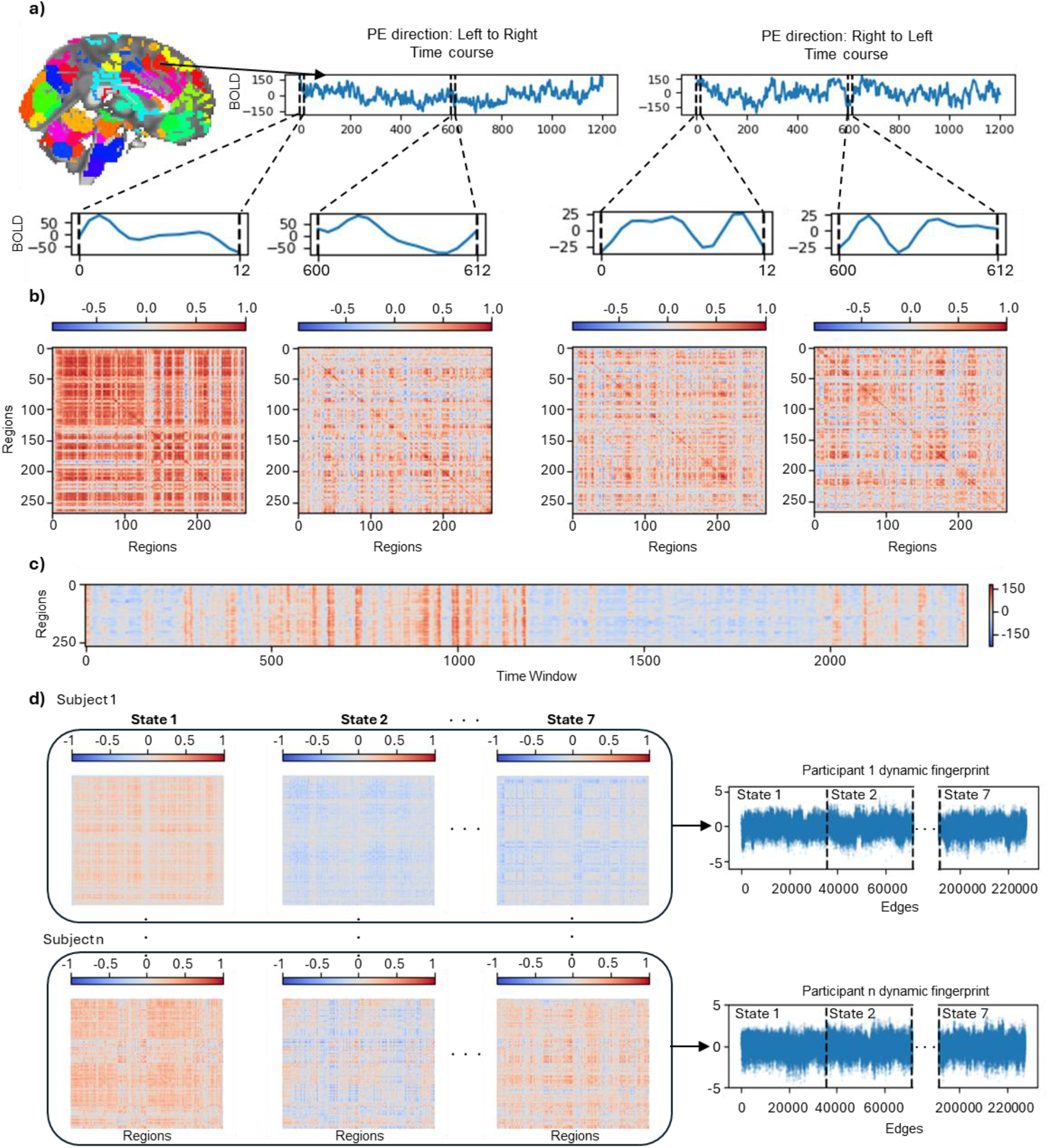
The pipeline for finding state-specific connectomes and dynamic fingerprints for the participants. (a) The time course of each ROI (top right) is obtained by averaging the signal of voxels within that ROI (top left). Using a sliding time window, we focus on different windows of the signal for the analysis (insets). (b) Functional connectomes computed by measuring the correlation between each pair of ROIs for the indicated windows in part (a). (c) Correlation strength vectors obtained by summing the correlation coefficients of each region within each time window. (d) Functional connectivity patterns of states for each participant. The lower triangle of the connectomes is taken and vectorized. The dynamic fingerprints are made of normalizing these vectors and concatenating them.

For each scan, the time series of each ROI was obtained by spatially averaging signal intensities of its voxels at each time point. The time series were preprocessed by linear detrending and regressing out motion parameters and their derivatives, as well as the average global signal. Additionally, a low-pass filter with a cutoff frequency of 0.25 Hz was applied.

For each phase-encoding direction of each scan, the time courses of regions were divided into different time windows (Figure 1.a) based on the desired window size (see section 2.4) of the analysis, and a window stride of 1 TR (0.72s). Next, Pearson’s correlation coefficient was calculated between each pair of ROIs to derive the functional connectome in each time window (Figure 1.b). The resultant functional connectivity maps were demeaned by subtracting the mean connectome across windows and participants; this was done separately for each day and each phase-encoding direction, respectively.

### 2.3 Obtaining the brain connectivity states

The average brain connectivity states across all participants were identified by using the following procedure. First, we summed the correlation coefficients of each ROI in each window to obtain the connectivity strength vectors for each participant’s connectome (Ou et al., 2013) (Figure 1.c). Then, we performed hierarchical clustering on the connectivity strength vectors of each participant, where the distance measure was the inverse of the correlation coefficient between two vectors and the number of clusters was set to 20 (Ou et al., 2013). This resulted in 20 cluster center vectors for each subject.

Using the same distance measure, group-level hierarchical clustering was applied to participant-level cluster centers of day 1 data to obtain the correlation strength cluster vectors of the seven brain states. Then, each participant’s connectivity strength vector for each window was correlated to the state cluster vectors, and each window was labeled as the state with the highest correlation coefficient. By averaging the functional connectivity maps of the windows that belong to the same state across the participants, the connectivity pattern of each state was obtained. Furthermore, participant-based connectome of each state was obtained by averaging the same-labelled connectomes for each participant (Figure 1.d).

### 2.4 Selecting the optimal sliding window size and obtaining the fingerprints

The size of the sliding window was initially set to the shortest length reported in the literature to capture the brain dynamics (8 seconds or 11 repetition times (TRs)) (Shakil et al., 2016). While various numbers of states have been used to describe the brain dynamics at rest (Chen et al., 2016, Hussain et al., 2023, Vidaurre et al., 2017), we did a preliminary analysis using the initial window size to identify the optimal number of states, and observed that for numbers more than 7, some participants did not go through all of the states, so 7 was set as the number of brain states. Using the resultant dynamic connectivity states of the preliminary analysis, the average duration of each state (number of TRs that a subject’s brain remained in a certain connectivity state) over their occurrences was calculated across the participants. The window size for the main analysis was chosen as the shortest average duration of states among all states in the preliminary analysis. This average duration was rounded down to the nearest integer to define the window size in terms of number of TRs.

With the optimal sliding window size set as described above, state-specific connectomes (i.e. the connectivity patterns) of each participant were found. State-based connectomes for each participant are symmetric 268 × 268 matrices and only the elements below the diagonal of these matrices were used. For each participant, the state-specific connectomes were vectorized and Fisher r-to-z transformed to obtain their state-specific fingerprint. The participants’ dynamic fingerprints were constructed by concatenating their state-specific fingerprints across all states (Figure 1.d).

### 2.5 Static fingerprints

Static fingerprints were formed using the procedure outlined in (Finn et al., 2015). Time courses of different regions were calculated for both phase encoding directions and concatenated. Then, pairwise correlation coefficient between different regions were calculated and z-score normalized to obtain the stationary connectivity matrix for each participant.

### 2.6 Identification based on dynamic and state-specific fingerprints

To evaluate the effectiveness of the dynamic fingerprints in individual identification, each participant’s fingerprint based on data on day 1 was set as the target and the Pearson’s correlation coefficient was computed between the target and day 2 fingerprints of all participants. If the highest correlation is found between the target and the same participant’s day 2 fingerprint, it indicates correct identification for that individual. The same process is applied for identifying day 2 target fingerprints based on day 1 data. The overall accuracy for each day is determined by the ratio of correct identification.

In addition, the performance of each single state in identification was examined by applying the same process on state-specific fingerprints. The identification accuracy of these states was compared to one another, as well as to the static case. This analysis identifies which states are most informative for individual identification and how their performance compares to using a stationary approach.

To assess the statistical significance of the identification results, we performed permutation tests for both state-specific and dynamic fingerprints identification accuracy. In either case, this was done once with day 1 participants as the database and shuffling the day 2 participants identities, and once with day 2 database with randomizing day 1 indices. Ten thousand permutations were conducted to obtain the null distribution of the accuracy happening by chance and the significance level considered was p-value below 0.001.

### 2.7 Edgewise contribution to identification using state-specific connectomes

Differential power (DP) (Finn et al., 2015) was calculated to examine which functional connections contribute significantly to individual identification. This parameter shows how powerful each functional connection (i.e. edge or feature) is in differentiation across the participants. We followed the steps described by (Finn et al., 2015). In particular, we computed an empirical probability for each edge that quantifies its strength in identification.

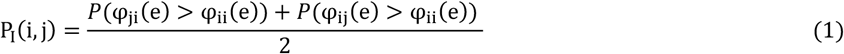

Here, i and j are labels of different participants (i≠j), e denotes a particular edge, φ_ij_(e) refers to the product of values of edge e between participants i and j (each from a different day), and N is the sample size. For an edge to be distinguishing between the participants, this product should be higher when calculated for the same participant than for different participants. Therefore, a smaller P_I_ is associated with a better differentiation power for the edge. The overall differential power of an edge is computed by determining the log likelihood of P_I_ across all participants.

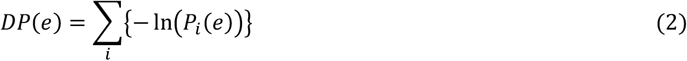

### 2.8 Cognitive behavioral score prediction

The ability of state-specific fingerprints to predict behavior was investigated and compared to that of static fingerprints. Scores from the Penn progressive matrices (PPM), Dimensional Change Card Sort (CS), Flanker task (FL), and Picture Vocabulary Test (PV) of the HCP data were used for this analysis (Cai et al., 2021). Each test assesses different aspects of cognition and recruits a certain functional domain of the brain for performing the task (Table 1). Prediction was performed using all states separately as well as the static fingerprints. For the prediction task, with both the state-specific and static approaches, we only used data from scan session 1 of the participants.

**Table 1.** Summary statistics and the functional domain of the cognitive tasks.

| Task | Functional Domain | Summary Statistics |  |
| --- | --- | --- | --- |
| | | Mean ± SD | Range |
| Penn Progressive Matrices (PPM) | Fluid Intelligence | 16.28 ± 4.29 | 5 - 24 |
| Dimensional Change Card Sort (CS) | Cognitive Flexibility/Execution | 101.81 ± 10.21 | 66 - 122 |
| Flanker Task (FL) | Cognitive Inhibition/Execution | 102.55 ± 9.58 | 73 - 123 |
| Picture Vocabulary Test (PV) | Language Comprehension | 105.97 ± 14.36 | 68 - 146 |

We used a leave-one-out (LOO) approach to predict the cognitive scores of each task with linear regression. In doing so, firstly a participant’s data was removed and the mean frame-to-frame displacement across the participants was regressed out of both the behavioral scores as well as each functional connection’s value (O’Connor et al., 2021). This was later applied to the test participant to remove the effects of average displacement from their data as well. Then, a feature selection step was performed on the training set, where the behavioral scores were correlated with each edge of the connectomes across the training set, and if the p-value of this correlation was less than the feature selection threshold (FST), that feature was retained for model building. These selected features were grouped based on the sign of their correlation (negative or positive) and used to build respective predictive models for each task. Then, correlation strength, derived by summation of the selected features values, was computed for each participant in both negative and positive models. Finally, these correlation strengths were used to develop a linear regression model to predict the behavioral scores of the training set and were applied to the left-out participant to predict their score. This process was done for all data points and a vector of the predicted scores was generated. Model performance was assessed by computing the correlation coefficient between the vector of predicted scores and the actual (observed) scores.

## 3 Results

### 3.1 States average duration

The average durations of the states using a window size of 11 TRs were computed for all of the states (Figure 2). Each state had a different average duration, where state 4, with the lowest average duration (12.1 TRs), determined the final window size of 12 TRs, ensuring the sliding window could capture state transitions.

**Figure 2.**
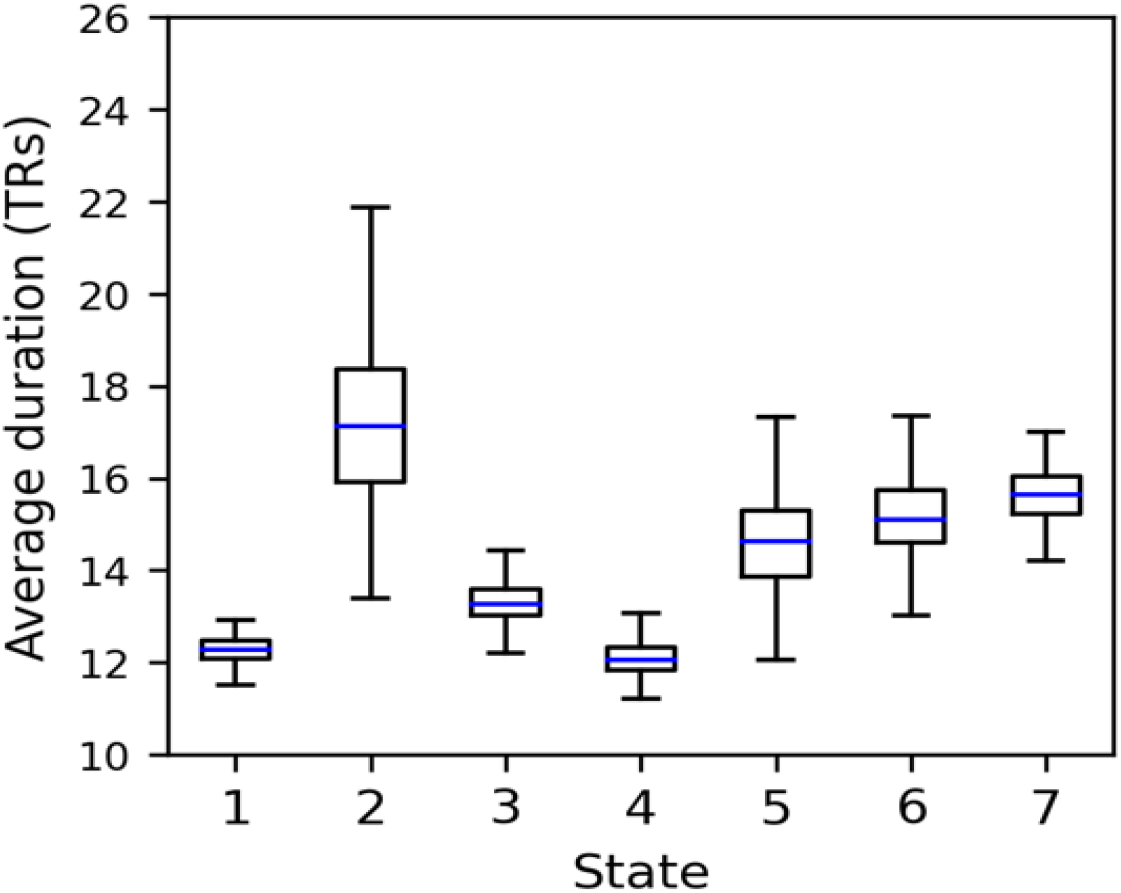
Duration of the states. The residence time of different states is summarized using quartiles, with the blue line indicating the average duration across participants. These are the average duration of different states given that the sliding window length was set to 11 TRs. The state with the shortest time span, which is state 4 in this figure, determines the length of the window needed to capture the dynamic transitions.

### 3.2 Brain states connectivity patterns

Using the pipeline outlined in 2.3, and by setting the window size to 12 TRs, the states connectivity patterns were obtained. Each of the brain states exhibits a distinct connectivity pattern that includes state-specific correlations within and between different known functional networks (Figure 3). The functional connectivity of the states shows a range of values, with some states dominated by positive or negative connectivity. This shows the ability of our state-based approach in capturing the dynamics of the brain.

**Figure 3.**
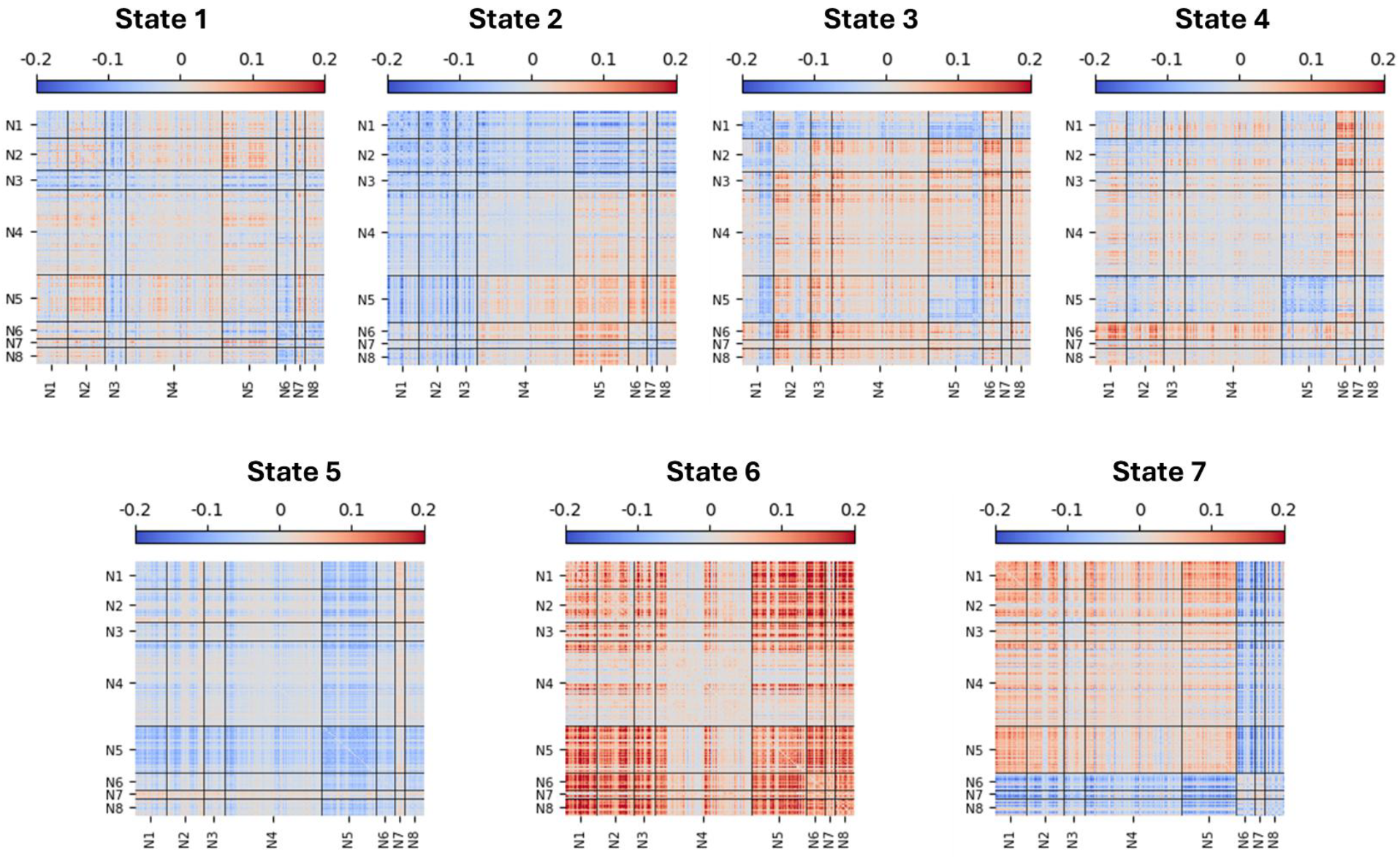
Connectivity patterns of the brain states. The ROIs have been relabeled to visualize patterns based on brain networks. The black lines separate the regions belonging to the same network from others. N1, the medial frontal network; N2, the frontoparietal network; N3, the default mode network; N4, the subcortical-cerebellum network; N5, the motor network; N6, the visual I network; N7, the visual II network; N8, the visual association network. Red elements show positive correlation between ROIs, while the blue ones indicate negative correlations.

### 3.3 Identification

The static fingerprints successfully identified 189 participants (87.91%) when day 2 was set as the target, and 195 (90.70%) when day 1 was the target. On the other hand, the accuracy of identification based on the dynamic fingerprints outperformed the static method in both days, with 97.21% and 95.81% correct identification rates, for day 1 and day 2, respectively.

Considering the fingerprints of each state separately, the states had varying rates of identification (Table 2). State 2 had the highest accuracy, with 95.81% and 95.35% of the participants identified correctly, for day 1 and day 2, respectively, showing the key role of this state in the distinguishing power of the dynamic fingerprints. States 5 and 7 also had high identification accuracy (80+%) on both days, which were close to the stationary fingerprints. State 6 had the lowest accuracy, where only 57.2% of day 1 targets, and 60.90% of day 2 targets were identified correctly. The permutation test showed that for both the dynamic and state-based fingerprints, the accuracy results were significantly above chance (p<10^−3^).

**Table 2.** Identification accuracy of the brain states. State-specific fingerprints performance in identification.

| State | Day 1 target (%) | Day 2 target (%) |
| --- | --- | --- |
| 1 | 59.53 | 63.72 |
| 2 | 95.81 | 95.35 |
| 3 | 77.67 | 79.53 |
| 4 | 75.81 | 73.95 |
| 5 | 90.70 | 89.30 |
| 6 | 57.21 | 60.93 |
| 7 | 80.00 | 85.58 |

### 3.4 Edgewise contribution to state-specific connectomes in identification

Each edge represents a functional connection between two different brain regions, and finding the edges that play a role in individual differentiation will aid in understanding individual differences in brain function and health. Thus, for each of the states as well as the static case, we calculated the DP of all edges and thresholded them at 99.9 percentile. Similar to (Finn et al., 2015), we computed the fractions of the supra-threshold DP edges between the functional networks and within each network (Figure 4).

**Figure 4.**
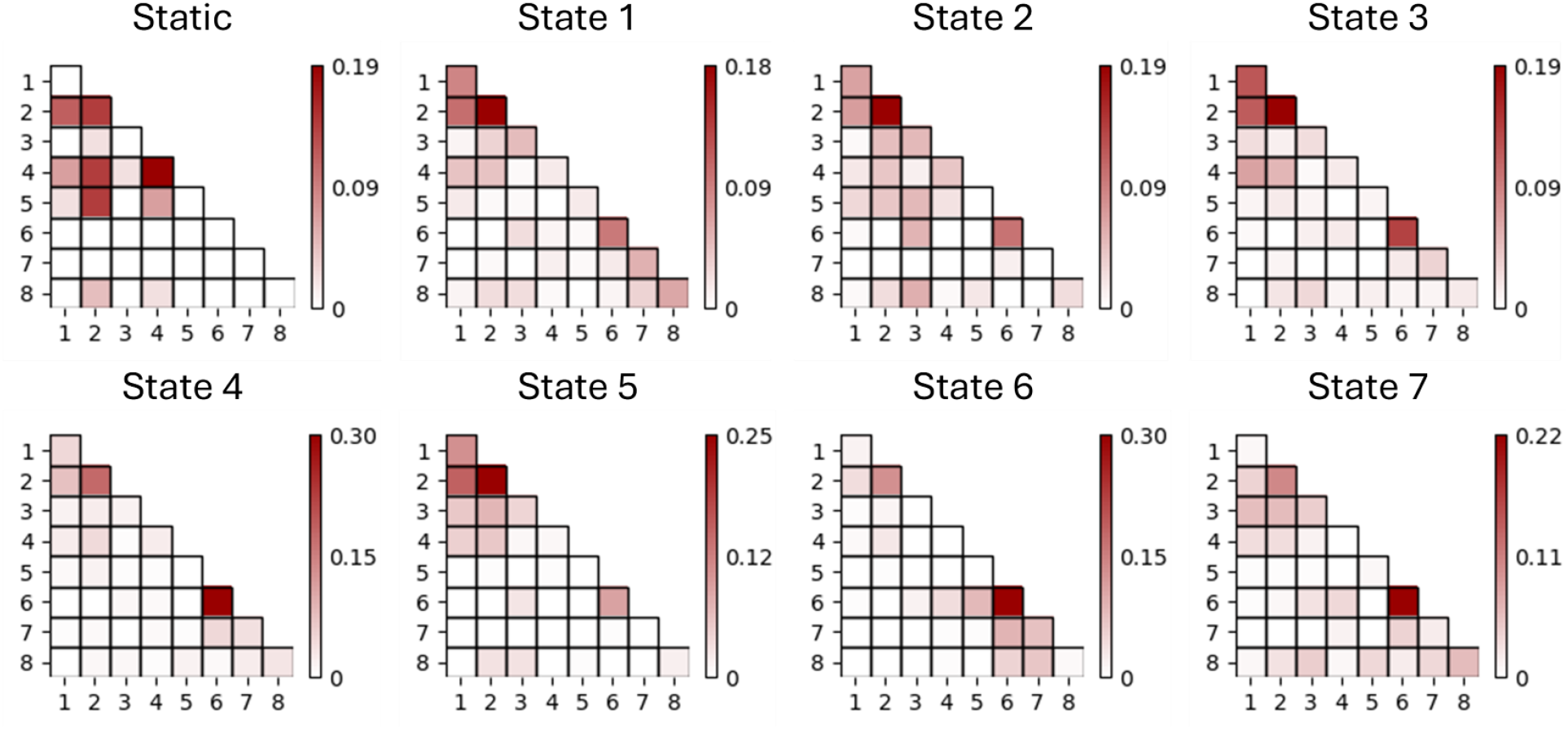
Edgewise contribution to identification performance of the states. The color bars show the fraction of edges with DP in 99.9 percentile. Each element in the grids is dedicated to connections within or between two certain networks. The list of networks is as follows: 1, the medial frontal network; 2, the frontoparietal network; 3, the default mode network; 4, the subcortical-cerebellum network; 5, the motor network; 6, the visual I network; 7, the visual II network; 8, the visual association network.

In the static case, the frontoparietal and the subcortical-cerebellum networks played a key role in differentiating between participants’ connectomes. The edges within both networks as well as the functional connections between them had the highest fractions of supra-threshold DP edges. The connections between the frontoparietal and the motor network also contributed to individual identification in the static case.

Looking at the state-specific fingerprints, we again observed that a large fraction of the supra-threshold edges was within the frontoparietal network, as seen in the static case. However, in the dynamic states, connections within the visual I network played a significant role in subject identification, particularly in states 4, 6, and 7—a contrast to the static case, where these edges were less prominent. Meanwhile, the subcortical-cerebellum network, which had a substantial contribution in the static case, played a less prominent role in any of the dynamic states.

### 3.5 Cognitive score prediction

Other works have shown that different measures of cognitive ability can be predicted using functional connectome (Finn et al., 2015, Lv et al., 2024, Manglani et al., 2022). Here, we compared the predictive power of the static and state-specific connectomes using the correlation between the predicted and observed scores, and reported its values for FSTs of 0.01, 0.05, and 0.1 with both the negative and positive models (Figure 5).

**Figure 5.**
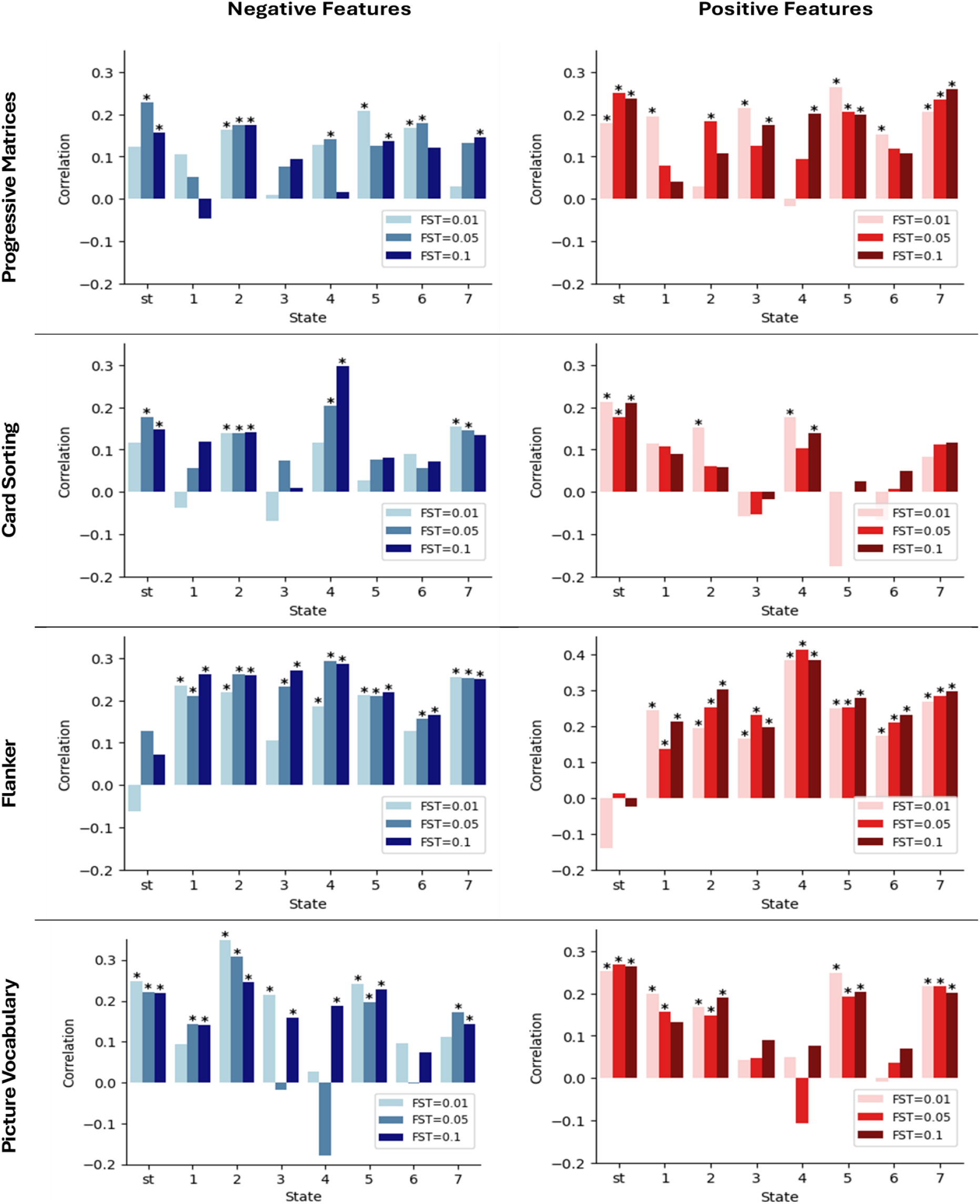
Cognitive tests score prediction by each state and static connectome. The bars show the correlation coefficient between the predicted and observed scores for each feature selection threshold (FST). On the x-axis, “st” represents the static connectomes, while the numbers refer to each state. The right column (charts with the reddish bars) shows models built with the positive features, and the left column represents negative models. The bars indicated with a * show a significant (p<0.05) correlation between the actual and predicted scores.

For PPM and CS tasks, the static connectome had a similar prediction performance, where it successfully predicted (all p-values < 0.05) the scores in both negative and positive models using all FST values except for the negative model using FST of 0.01. From a dynamic point of view, state 5 and state 4 connectomes were the most successful in predicting the PPM and CS scores, respectively, both producing the most accurate prediction of their corresponding task scores across all models. Using a positive model and an FST of 0.01, state 5 connectome outperformed all other dynamic models as well as static connectome in predicting the PPM scores (r=0.26, p<10^−4^). State 4 negative features exhibited the best prediction of CS, where the model predicted this task’s scores with a correlation coefficient of 0.3 using an FST of 0.1 (p<10^−4^).

In predicting the FL scores, the static connectome showed no predictive power. In both negative and positive models, none of the static regressors predicted the scores of the FL task with significant correlations, whereas almost all of the dynamic states performed well using all FSTs. The highest correlation across both models was for state 4 positive model using an FST of 0.05, where the correlation between the observed and predicted FL scores was r=0.41 (p<10^−6^). State 4 with an FST of 0.05 was again the most predictive of the FL scores among all negative models (r=0.29, p<10^−4^).

For PV scores, the static connectome, state 2 connectome, and state 5 connectome predicted the task results successfully, and the highest score observed was state 2 negative model with FST set to 0.01, where a correlation of r=0.35 was reported between the observed and predicted scores (p<10^−6^).

For each task, we examined the selected features in the dynamic states and the static connectome to ascertain which brain networks contributed to prediction of each task. We first chose the most successful dynamic state for each task by selecting the state with the highest average correlation with the observed scores across all models. Following this procedure, states 5, 4, 4, and 2 were chosen for PPM, CS, FL, and PV tasks, respectively. For better visualization, we only showed functional connections whose correlation p-value with the cognitive scores were below 5×10^−4^ for all the folds of the LOO approach (Figure 6).

**Figure 6.**
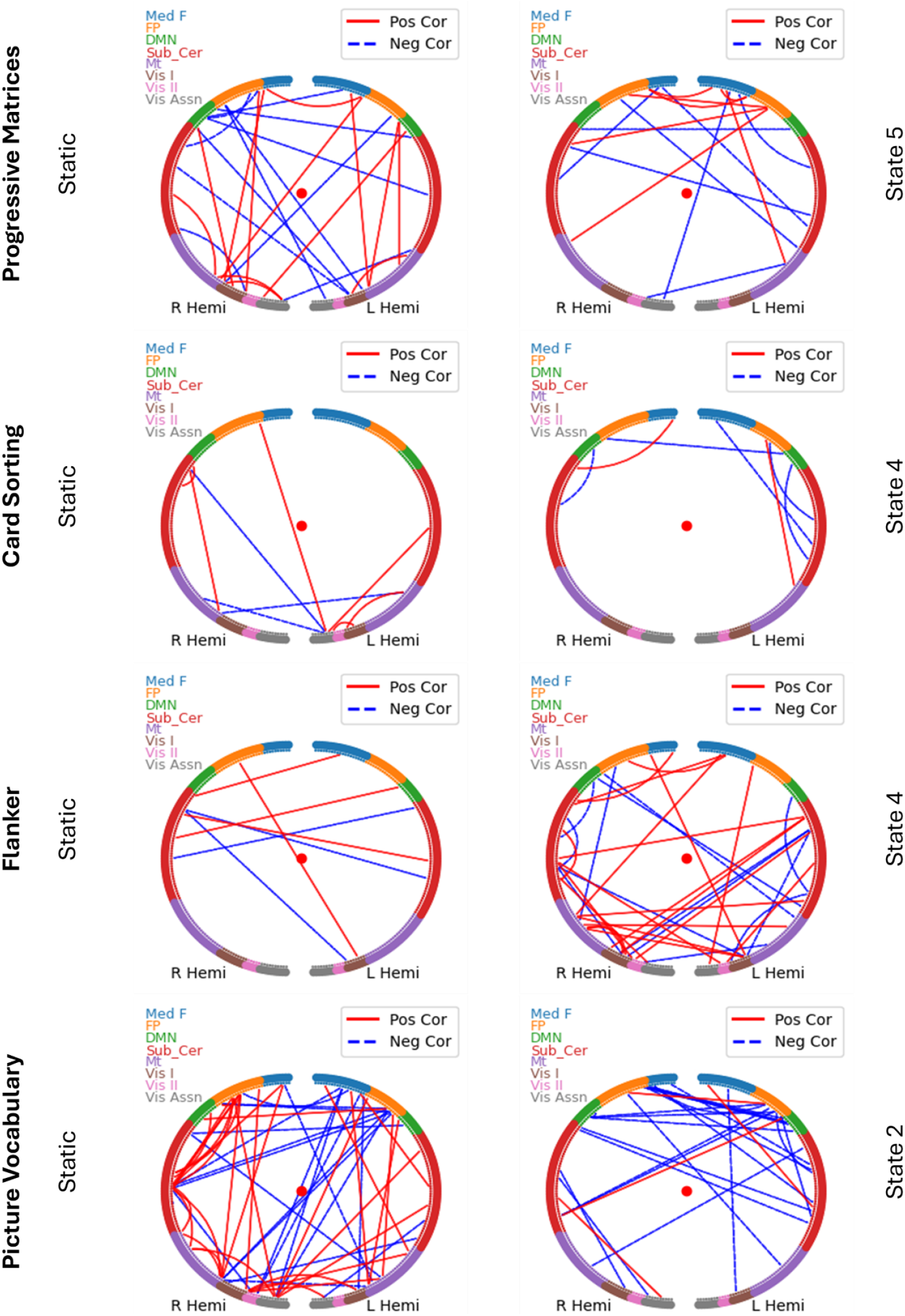
The features predictive of the cognitive scores. The edges (functional connections) that had a correlation p-value less than 5×10^−4^ in all folds during the feature selection process with the scores of each task are visualized. The blue lines show features negatively correlated with the scores, while the red ones are showing positively correlated ones. The graphs on the left belong to the static connectomes, and the ones on the right show the most predictive dynamic state of each of the tasks. L hemi and R hemi represent left and right hemispheres, respectively. The network abbreviations are: Med F, the medial frontal network; FP, the frontoparietal network; DMN, the default mode network; Sub_Cer, the subcortical-cerebellum network; Mt, the motor network; Vis I, the visual I network; Vis II, the visual II network; Vis Assn, the visual association network.

With the PPM task, in the static model, the medial frontal networks of both hemispheres had positive connections with other networks and each other. The right primary visual network showed several positive edges in the static model but was less prominent in the dynamic model. On the other hand, the dynamic model (state 5) featured many positive connections involving the left frontoparietal network. For negative edges, the static model highlighted the right default mode, frontoparietal, and left primary visual networks, while the left subcortical-cerebellum network was the main contributor in the dynamic model.

For the CS task, fewer edges passed the threshold in either the static or the dynamic model compared to the PPM task. In the static model, approximately half of the supra-threshold positive and negative edges were connected to regions in the left visual association network. In the dynamic model (state 4), predictive edges were predominantly negative, involving the left subcortical-cerebellum network. Unlike the static model, frontal networks were more engaged in the dynamic model, while visual-related networks did not contribute to prediction.

With the FL task, very few features were predictive in the static connectome, and most of these features connected the right subcortical-cerebellum network to various regions in the left hemisphere. In contrast, for connectome of state 4, there were several negative and positive connections among all networks across the two hemispheres. Connections between the right primary visual network and the left subcortical-cerebellum network in state 4 were predictive of FL. The right motor network of state 4 also had a highly pronounced role in providing positive edges correlated with the FL scores.

For the PV task, the static model showed a high concentration of positive connections between the right subcortical-cerebellum and right frontoparietal networks, while the left frontoparietal network had negative connections with several right-hemisphere regions. In the dynamic model (state 5), predictive connections were dominated by negative edges, primarily linking the left default mode network to the right medial frontal and frontoparietal networks. Negative predictive features of this state also involved the right default mode and left frontoparietal networks, the latter of which was similar to the static model.

## 4 Discussion

Recent research highlighted the ability of functional connectivity profiles to be used as a unique identifier, similar to a fingerprint, and distinguish individuals in a large population (Finn et al., 2015, Hu et al., 2022, St-Onge et al., 2023). These studies used static functional connectivity to generate participant-specific connectome-based fingerprints for identification. However, the brain alternates between varying states of connectivity in time and these temporal dynamics are participant-specific (Menara et al., 2021, Moretto et al., 2023). With seven connectivity states identified using a sliding time window and hierarchical clustering, we demonstrated that the identification accuracy of the dynamic fingerprint outperforms the stationary approach. The reduced identification accuracy of the static fingerprinting approach may be due to the absence of dynamic information in the static connectome because such dynamic information can be participant-specific and provide increased identification accuracy.

Each brain state is comprised of different patterns of connectivity within and between different networks. With this in mind, we also examined the identification accuracy of each brain state as the brain could have distinct within- and between-participant variability during each of these states (Finn et al., 2017). We observed that the most differentiating state connectivity pattern (state 2) was general negative correlation or non-correlation of the medial frontal, frontoparietal, and the default mode networks to other networks, while there was a positive connectivity within the motor network. The second differentiating state (state 5) had a general negative correlation between all networks. Prior work found that the frontoparietal network plays the most important role in distinguishing between different participants (Finn et al., 2015). Here, the states with a negative or near-zero correlation between this network and other networks show increased distinguishing power, indicating that the less correlated this network is with other networks, the more distinguishing the state becomes.

We also examined which brain networks contributed most to differentiation in the state-specific fingerprints by identifying the networks connected by edges with high differential power. We saw that in all states, a large portion of the connections were located within the frontoparietal network, in agreement with earlier work that investigated inter-subject variability (Cai et al., 2021, Cole et al., 2013, Finn et al., 2015, Finn and Todd Constable, 2016, Gao et al., 2014). The nodes within these networks are located in higher-order brain systems with inter-subject variability seen in their functional connectivity profiles (Wang et al., 2021a), making them the leading network in providing identifiability for the functional connectome (Colenbier et al., 2023). We also saw the connections within the visual I network contributed significantly to identification, which indicates the difference that the visual perception and connections associated with visual processing makes among individuals (Zhao et al., 2021). This is in agreement with the notion that at shorter temporal windows, functional connections in visual networks have a more emphasized role in fingerprinting of the connectome (Van De Ville et al., 2021), and that is why their role in identification is prominent in state-specific fingerprints, but not in the static case.

We revealed that certain states effectively predicted measures of intelligence, cognitive flexibility, inhibition, execution, and language comprehension. The performance of these states varied depending on the functional domain of the cognitive and behavioral tests, as well as the feature selection threshold. Notably, at least one of the dynamic states outperformed the static connectome in each of the tasks, highlighting how dynamic information within these states enhances the connectome’s ability to describe cognitive measures. This demonstrates the advantage of dynamic connectivity states in capturing individual differences in cognition and suggests their potential to be used as neurocognitive biomarkers.

States 2, 4, and 5 predicted all four cognitive task scores for most of the feature selection thresholds, indicating that these brain states are more informative compared to other brain states. State 4 showed high connectivity of the primary visual network. This is in correspondence with earlier work (Santarnecchi et al., 2017), where it was shown that in processing a task at the stage of applying a learned rule to novel stimuli (rule application), the primary visual network is at a high connectivity state. There is also high connectivity of the motor network with itself and the visual-processing networks in the connectome of state 2. These perceptual and sensory networks move into a state of alertness with high connectivity, which is essential for performing a cognitive task, making the states with such connectivity patterns the most predictive ones. On the other hand, the frontal high-order networks are in a state of negative or no correlation in these predictive states, which helps them preserve their original shape and contribute to describing individual differences in cognition. Conversely, in the states where the frontal networks are being positively correlated (synchronized) with other networks, the states were not informative of the cognitive abilities of the individuals, and poor prediction performances were seen (states 1, 4, 5, and 7).

The implication of various brain networks can be seen by analyzing the contributing features to the best predictor state of each task. There was a large contribution of the medial frontal and the frontoparietal network in fluid intelligence (PPM task) prediction, consistent with previous findings in both dynamic and static models (Cole et al., 2013, Finn et al., 2015). For the CS task (executive function), few features passed the 5×10^−4^ threshold for both models, and contribution patterns of the two models were different. While most of the static predictive features were concentrated in the motor or visual networks, the majority of state 4’s predictive features were in the medial frontal network, frontoparietal network, and subcortical regions, consistent with findings of previous work (Cai et al., 2021). Looking at the Flanker task (inhibition function), a significantly higher number of predictive features was seen in state 4 connectome compared to the static method, possibly resulting in the superiority of the dynamic approach for prediction of this task. The role of the default mode network and the subcortical-cerebellum network in the prediction of this score in state 4 connectome was more prominent than other networks. This is consistent with previous reports that the subcortical brain regions in these networks connect with various cortical and subcortical brain regions that are effective in terms of processing the visual-spatial information (Molenberghs et al., 2007), aiding in performing well in an attentional control task such as Flanker. The default mode network also plays a role in external tasks by coupling with different brain networks and has increased connectivity during task states (Seeley et al., 2007). Finally, for the Picture Vocabulary task (language comprehension), the static connectome employed several connections within and between different visual networks to predict the scores, consistent with previous literature (Mantwill et al., 2022). This pattern was totally different in predictive features of state 2 connectome, where, similar to that for CS, the frontal and subcortical regions known to govern higher-order functions of the brain were the most prominent.

Several aspects of this work could be expanded in future research, building on the current findings. Finn et al. found the connectome derived from the medial frontal and frontoparietal networks had higher identification accuracy as compared to the whole-brain connectome (Finn et al., 2015). Future research could apply our dynamic fingerprinting approach on connectomes of these two networks and assess whether state-specific patterns derived from these networks enhance identification accuracy and see if the states have a better performance in predicting factors like aging and cognitive abilities compared to the current approaches (Loaliyan and Ver Steeg, 2024). Besides, parcellations other than the 268-node parcellation described by (Shen et al., 2013) could be used to derive the brain connectome, and the impact of parcellation on the performance of identification and prediction of dynamic fingerprints can be examined. Finally, we used a constant-length sliding time window and clustering to find the brain states. Other methods like the Hidden Markov Model (Chen et al., 2016) that do not inherently require such a window could be used to obtain brain states.

## 5 Conclusions

Our findings show that dynamic state-based connectomes, derived from resting-state fMRI data, could be used to identify individual participants and predict their cognitive performance. These states could potentially be used by neuroimaging studies as a basis for early diagnosis of certain cognitive diseases.

## Authorship contribution statement

**Amin Ghaffari:** writing – original draft, writing – review and editing, investigation, software, methodology, formal analysis, validation and visualization (lead); conceptualization (supporting). **Yufei Zhao:** writing - review and editing, methodology (supporting). **Xu Chen:** writing - review and editing, investigation (supporting). **Jason Langley:** writing - review and editing, conceptualization, investigation, and supervision. (supporting). **Xiaoping Hu:** writing - review and editing, investigation (supporting); conceptualization and supervision (lead).

## Authors’ disclosure statement

The authors declare no conflicts of interest related to this work. No competing financial or non-financial interests influenced the design, execution, analysis, or interpretation of the study.

## Funding statement

This research was conducted without any external funding or financial support

## Data availability statement

The data are publicly available through the HCP database (https://www.humanconnectome.org/) and were accessed in accordance with HCP data use terms and conditions.

## Notes

https://www.humanconnectome.org/

### Competing Interest Statement

The authors have declared no competing interest.

